# A Bayesian optimisation approach for rapidly mapping residual network function in stroke

**DOI:** 10.1101/2020.07.03.186197

**Authors:** Romy Lorenz, Michelle Johal, Frederic Dick, Adam Hampshire, Robert Leech, Fatemeh Geranmayeh

## Abstract

Post-stroke cognitive and linguistic impairments are debilitating conditions, with current therapies only showing small improvements. Domain-general brain networks seem to play a critical role in stroke recovery and characterising their residual function with functional neuroimaging (fMRI) has the potential to yield biomarkers capable of guiding patient-specific rehabilitation. However, this is currently challenging in patients as such detailed characterisation requires too many different cognitive tasks. Here, we use neuroadaptive Bayesian optimisation to overcome this problem, an approach combining real-time fMRI with machine-learning. By intelligently searching across many tasks, this approach rapidly maps out patient-specific profiles of residual domain-general network function. Whereas controls have highly similar profiles, patients show idiosyncratic profiles of network abnormalities that are associated with behavioural performance. This approach can be extended to diverse brain networks and combined with brain stimulation or other therapeutics, thereby opening new avenues for precision medicine targeting diverse neurological and psychiatric conditions.

## Introduction

Cognitive and linguistic impairments following brain injury such as stroke are a leading cause of disability, affecting over a quarter of a million people in the UK (Stroke Association UK) and over a million in the US (American Speech-Language-Hearing Association), with numbers expected to increase dramatically given the ageing population (Béjot et al., 2016). Current therapeutic strategies are only of limited success (Brady et al., 2016; Elsner et al., 2019; Merriman et al., 2019); therefore, there is a need for developing biomarkers that guide clinical prognosis as well as rehabilitation strategies. Given the great heterogeneity in stroke patients with respect to their pathophysiology and resulting functional deficits, functional magnetic resonance imaging (fMRI) is a promising method for discovering candidate biomarkers capable of distinguishing patient subgroups as it allows non-invasive mapping of brain (dys-)function with spatial precision. However, to date, no fMRI-derived biomarker is considered as ready to be used in clinical trials for predicting recovery of cognitive or language function (Boyd et al., 2017).

Nonetheless, fMRI measures of brain network activation and connectivity during task execution (“task-based fMRI”) show promising potential as clinically relevant biomarkers and thereby represent a developmental priority (Boyd et al., 2017). A major challenge for any progress in this direction is selecting the optimal task (or battery of tasks) to be administered to patients in the MR scanner.

This is because neither cognitive nor language-related functions can be readily mapped to distinct, single brain regions but rather emerge through the interaction between domain-specific (e.g., motor, auditory, language networks) and domain-general brain systems. Highly domain-general brain networks such as frontoparietal networks (FPNs) support processes including attention, working memory and learning (or reacquisition) of a skill (Duncan, 2010; Duncan & Owen, 2000; Fedorenko et al., 2013; Soreq et al., 2019). Damage to domain-general brain networks may explain why cognitive impairments seen in stroke patients are distributed across many different cognitive processes (Sachdev et al., 2004; Vasquez & Zakzanis, 2015). We have previously shown that intact domain-general brain regions are critical in recovery of language function following aphasic stroke (Brownsett et al., 2014; Geranmayeh et al., 2014, 2016, 2017) in keeping with studies confirming their role in recovery of motor deficits (Rinne et al., 2018) and the learning of pseudo language (Sliwinska et al., 2017). This builds a convincing case for their potential as a prognostic biomarker.

However, characterising residual function of domain-general networks in stroke patients is challenging because there is not a single, optimal task that is unique to probe each network; instead it involves quantifying network activation across many different cognitive tasks. However, such prolonged, multi-task neuroimaging protocols (King et al., 2019; Nakai & Nishimoto, 2020; Pinho et al., 2018) are practically unfeasible in patients. Meta-analytic approaches (de la Vega et al., 2018; Laird et al., 2009) are an alternative approach, but do not account for individual variability. Thus, the current status quo for clinical neuroimaging studies typically involves selecting a specific task (or small subset of tasks) in a relatively ad hoc manner, in the hope that it will reveal a useful neuroimaging biomarker. Given both the sheer volume of possible tasks and the constraints on scanner and patient time, this approach is unsuitable. Advancing task-based fMRI biomarker discovery requires a paradigm shift in order to be able to swiftly characterise residual brain network activity in individual patients using a diverse range of cognitive tasks.

Development of real-time analysis of functional neuroimaging data in combination with state-of-the-art machine-learning techniques (i.e., Bayesian optimisation) (Lorenz, Monti, Violante, et al., 2016; Lorenz et al., 2017) provides an unprecedented opportunity to derive subject-specific profiles of brain network function across multiple tasks in a short period of time (Lorenz et al., 2018), making it feasible to use in patients. Neuroadaptive Bayesian optimisation can efficiently search a large task space to identify the optimal set of cognitive tasks that maximise a predefined target brain network state in each individual (Fig. 1b). The approach’s efficiency stems from the intelligent search procedure: based on real-time analysis of the fMRI data, the machine-learning algorithm decides which task to test next in that particular subject. This is substantially faster than exhaustively testing all possible tasks while far more informative than selecting tasks at random.

**Figure 1:**
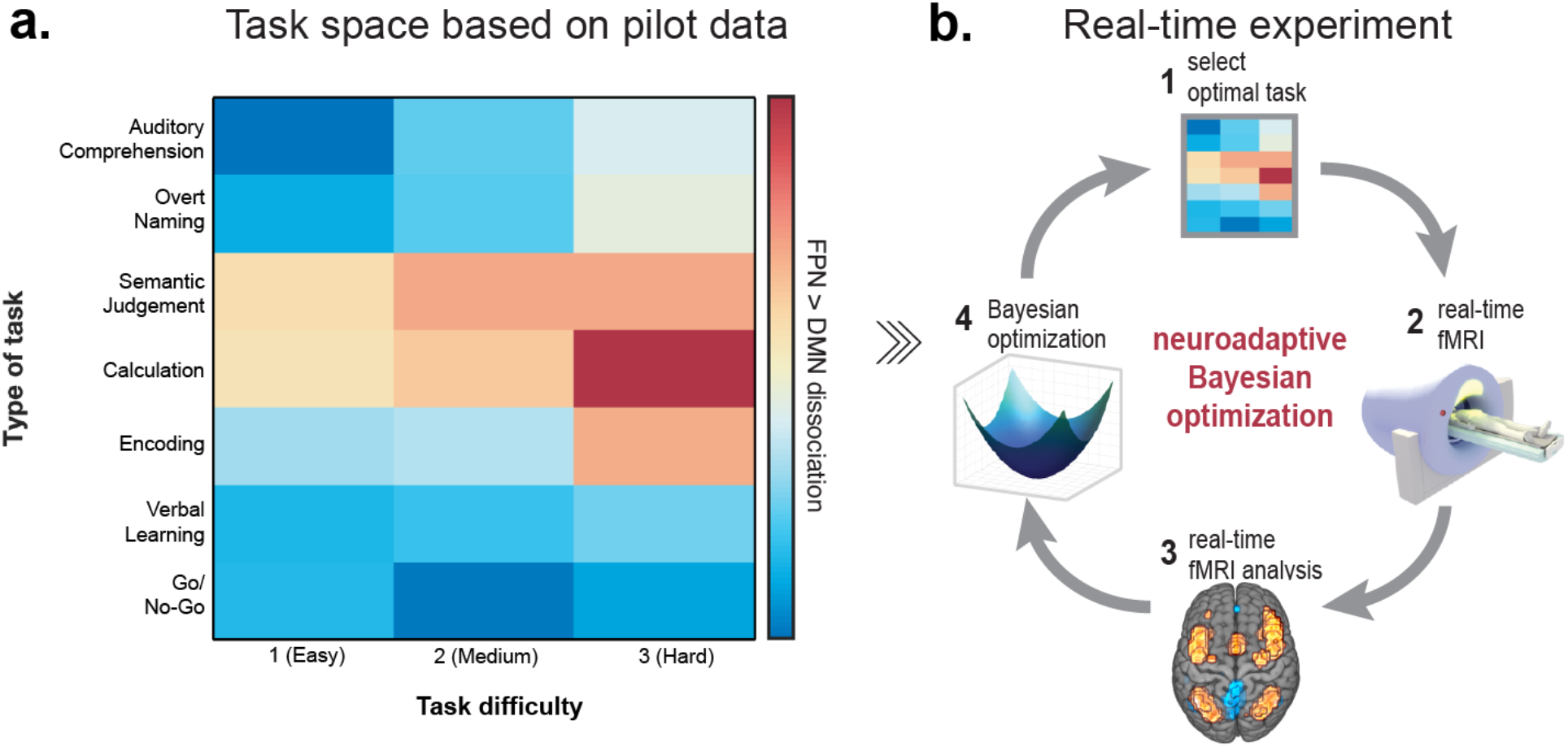
Overview of methodology. **(a)** We designed a two-dimensional (7×3) task space with one dimension corresponding to type of task, and the other to task difficulty. Tasks were ordered using pilot data collected separately in healthy volunteers. **(b)** This task space was searched through in our real-time experiment using neuroadaptive Bayesian optimisation. The aim was to quickly identify a subject-specific set of tasks that maximise the difference in activity between the FPN from the DMN. The method operates in the following steps: *(1)* the algorithm chooses a specific task x difficulty combination from the task space; *(2)* fMRI data is collected while the subject is performing the task; (*3)* the difference in brain level activation between FPN (warm colours) and DMN (cold colours) is computed in real-time; *(4)* the result from step 3 is used to update the algorithm (i.e., Gaussian process regression, see main text and Methods) and subsequently choose the next task x difficulty combination to be presented to the subject in the next iteration (back to step *(1)* in a closed-loop fashion).

Here, we apply neuroadaptive Bayesian optimisation to a cohort of left-hemispheric stroke patients with chronic aphasia (*n*=11) and demonstrate the approach’s potential for assessing patient-specific residual brain network function effectively and rapidly. For comparison, the method was also run in age-matched, healthy controls (*n*=14). Specifically, for each participant, we identify the set of cognitive tasks that maximally dissociate two domain-general networks, namely increasing activation in the bilateral FPN, and decreasing activation in the default-mode network (DMN). The choice of this target brain state was motivated by evidence suggesting that the difference in activity between the FPN and DMN was associated with language performance in left-hemispheric, aphasic stroke patients (Geranmayeh et al., 2016). Our results demonstrate that neuroadaptive Bayesian optimisation is a feasible and reliable neuroimaging technique for patient populations. In contrast to controls, who show highly consistent functional profiles of the FPN-DMN dissociation, patients’ profiles differ greatly, and exhibit unique profiles of network dysfunction that can be associated with out-of-scanner behavioural performance.

## Results

### Neuroadaptive Bayesian optimisation and task space design

Neuroadaptive Bayesian optimisation is substantially more efficient than randomly or exhaustively sampling all tasks because of two desirable properties: (1) it incorporates prior information about how the cognitive tasks relate to each other; and (2) guides its own sampling trajectory across tasks in an intelligent manner. The intuition behind (1) is that tasks that are expected to elicit a similar brain response are grouped together in the search space, while dissimilar tasks are grouped further apart. Thanks to this prior information, the algorithm does not need to test all possible tasks in the real-time optimisation run, but instead can sample a few, highly informative tasks and then make predictions for all other tasks by applying a nonlinear spatial regression (i.e., Gaussian process (GP) regression). This allows the algorithm to swiftly identify regions in the search space (i.e., a set of tasks) that are suboptimal for its optimisation aim (i.e., maximising FPN-DMN dissociation) and instead focus on sampling tasks from the optimal regions in the search space.

This highlights the importance of appropriately designing the task space for the algorithm to search through. Here, we designed a two-dimensional task space with one dimension corresponding to *type of task* and the other to *task difficulty*. We selected three cognitive tasks (Calculation, Go/NoGo and Encoding) and four language tasks (Overt Naming, Auditory Comprehension, Semantic Judgement and Verbal Learning). Tasks were chosen according to three criteria: (1) their ability to assess core cognitive and language deficits, (2) their predicted probability of recruiting the FPN (Yeo et al., 2014) and (3) the ability for patients to perform and understand these tasks (Price & Friston, 1999). Whereas in our past work, we have aligned tasks in the search space based on a previous meta-analysis (Lorenz et al., 2018), here we added three tasks (Auditory Comprehension, Semantic Judgement and Verbal Learning) that were not part of this meta-analysis. Therefore, to order these seven tasks along the task dimension, we used pilot data collected prior to the real-time study in 8 healthy volunteers (3 female, mean age ± SD: 27.9 ± 8.3 years). Each task had three levels of difficulty with increasing complexity and cognitive demand, resulting in a total of 21 different task x difficulty conditions the algorithm could choose from.

For the real-time study, each participant underwent two, independent optimisation runs, for which the algorithm was re-initialised and thus blind to any data collected in the subject’s previous run or any previous subjects. This resulted in different search trajectories of the algorithm through the task space for each run and participant, and therefore allowed us to assess the reliability of each participant’s results.

### Most patients are able to perform multiple tasks in the scanner

14 aphasic stroke patients (mean age ± SD: 58.57 ± 10.43 years, mean post stroke time ± SD: 5.52 ± 3.25 years) participated in our study. For 5 of 14 patients, task instructions had to be provided orally via a microphone due to reading impairments. For 3 patients (030, 032 and 040) the second run was discarded due to distress and/or fatigue, causing us to stop the run prematurely. These 3 patients are excluded from all analyses as we could not guarantee the validity of the patient-specific results. In addition, from the 15 controls (mean age ± SD: 56.73 ± 6.76 years) that were scanned, one had to be excluded as auditory stimuli could not be heard due to a technical issue. Thus, all analyses are based on 11 patients (4 female, mean age ± SD: 59 ± 10.9 years, mean post stroke time ± SD: 5.95 ± 3.42 years) and 14 controls (8 female, mean age ± SD: 55.6 ± 6.8 years).

Given the nature of the clinical populations, we first assessed whether patients performed above chance while undergoing the scan, indicating their understanding of the various task instructions. For this, non-parametric effect size measures (i.e., area under the receiver operating characteristic curve, AUROC) were computed for each task condition separately, comparing patient group-level accuracy with chance level. Significance was determined when the one-sided lower 95% confidence bound was higher than an AUROC of 0.5. This analysis could not be performed for the easiest level of the Auditory Comprehension task (as all trials were correct for this difficulty level) and the Naming task (no chance level could be determined as this task did not require button presses, see Methods). Results (Fig. 2a - left panel, corresponding lower confidence bound is listed in Table 1) demonstrate that patients performed above chance for all difficulty levels of the Go/No-Go task, for the easiest and medium levels of the Calculation task and the easiest level of the Semantic Judgement task. Further, they performed above chance for the most difficult level of the Auditory Comprehension task but not for the medium level – which can likely be explained due to unequal sampling across both conditions (i.e., the medium level was only sampled 3 compared to 18 times of the most difficult level, see Fig. 2d for sampling frequency/sample size ratio). With respect to the Encoding task, patients performed only above chance for the medium difficulty level. Given that the lower confidence bound for the easiest level of the Encoding Task is 0.4903 (Table 1) and AUROC values are > 0.6 for the easiest and medium level of this task, it can be assumed that patients performed higher than chance – even though it appears that this task is among the harder tasks we tested. Patients did not perform above chance for any level of the Verbal Learning task. When comparing this to controls (Fig. 2a - middle panel), we see that controls did not perform above chance for Verbal Learning Task either, illustrating that this particular task was ill-designed (see Methods). The largest performance differences between patient and controls were found for the most difficult conditions of the Semantic Judgement, Calculation and Encoding tasks, followed by medium levels of the Auditory Comprehension and Semantic Judgment tasks (Fig. 2a – right panel).

**Table 1:** One-sided 95% lower confidence bound for AUROC.

| Task | Patients |  |  | Controls |  |  |
| --- | --- | --- | --- | --- | --- | --- |
|  | Difficulty 1 | Difficulty 2 | Difficulty 3 | Difficulty 1 | Difficulty 2 | Difficulty 3 |
| Auditory Comprehension | Nan | .4059 | .7015 | Nan | .9935 | .664 |
| Naming | Nan | Nan | Nan | Nan | Nan | Nan |
| Semantic Judgement | .6464 | .4407 | .4741 | .9979 | .9993 | .8920 |
| Calculation | .7250 | .7884 | .4011 | .9946 | .7802 | .8813 |
| Encoding | .4903 | .5026 | .3321 | .7971 | .5946 | .7168 |
| Verbal Learning | .2500 | .3134 | .4229 | .4057 | .3731 | .3322 |
| Go/No-Go | .9448 | .9982 | .9899 | .6707 | .9971 | .9030 |

**Figure 2:**
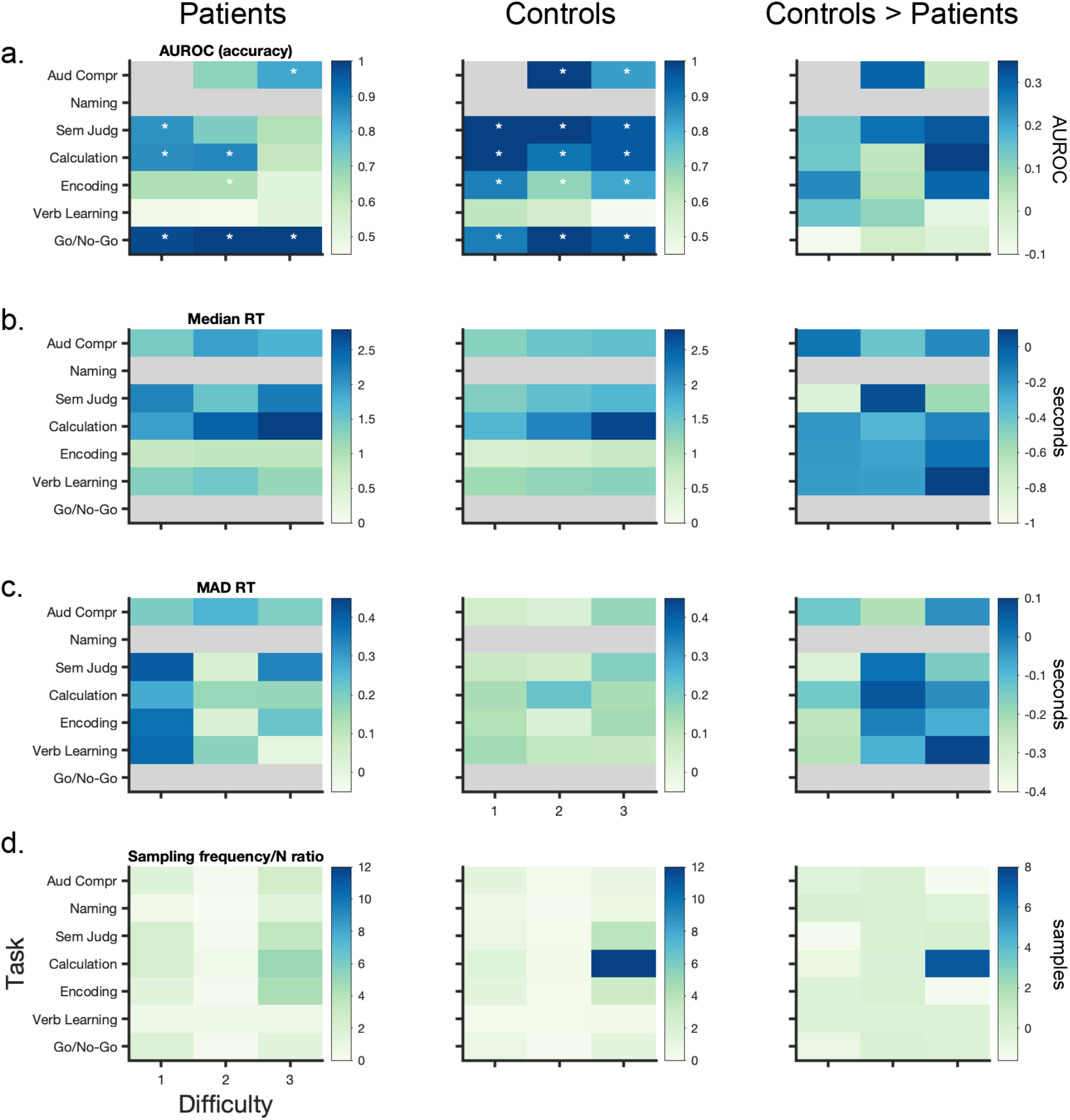
Behavioural results and sampling behaviour of algorithm. All results are listed for patients (left panel), controls (middle panel) and the difference between controls and patients (right panel). **(a)** The area under the receiver operating characteristic curve (AUROC) is a non-parametric effect size measure indicating the difference between the empirically obtained accuracy and chance level for each task. Stars indicate the tasks for which patients and controls performed significantly above chance; significance was determined using a one-sided 95% lower confidence bound criterium of AUROC > 0.5, exact values are listed in Table 1). **(b)** Median and **(c)** variance of reaction time (i.e., median absolute deviation (MAD)) across task space. **(d)** Given the algorithm’s subject-specific trajectories through the task space, each subject is exposed to a different set of tasks; here, we show the absolute number of times each task was selected by the algorithm – corrected for the difference in sample size N across both groups (i.e., N=11 for patients and N=14 for controls). Grey shaded areas correspond to NaN: accuracy could not be computed for the Naming task (as no chance level could be determined) and the first difficulty level of the Auditory Comprehension (as it was always correct); reaction time could not be computed the Naming task (as no button press was required) and for Go/No-Go tasks (as subjects were instructed to inhibit a response).

### Patients perform less accurately, slower, and more variably than controls

To assess behavioural performance, fMRI measures, in-scanner motion and the relationship between fMRI and behaviour, linear mixed-effect models (LME) were computed (all models are listed and numbered in Table 3 in the Methods sections; we refer to the corresponding number of each model in the following sections as LME X.X). Median reaction times (RT) and median absolute deviation (MAD) of reaction times for each task are shown in Fig. 2b and Fig. 2c, respectively. As expected, overall patients performed less accurately (LME A.1: Group *t*(746) = −4.13, *p* < .001) and slower than controls (LME A.2: Group *t*(643) = 5.53, *p* < .001). Both patients and controls performed less accurately (LME A.1: Difficulty *t*(746) = −4.76, *p* <.001) and more slowly for more difficult task levels (LME A.2: Difficulty *t*(643) = 9.84, *p* < .001). Patients’ accuracy was not differentially affected by task difficulty compared to controls (i.e., no significant interaction effect, LME A.1: Group x Difficulty *t*(746) = −0.95, p < .034); in fact, they showed a gentler increment in response time with increasing difficulty compared to controls (LME A.2: Group x Difficulty *t*(643) = −2.58, *p* = .01). This is due to patients’ considerably slower responses for the easiest task conditions relative to controls (Fig. 2b) and that there was a set time window to respond for each task (i.e., tasks were not self-paced; see Methods). Whereas reaction times decreased from the first to second run in both groups (LME A.2: Run *t*(643) = −2.98, *p* = .003), only patients were more accurate in the second run (LME A.1: Group x Run *t*(746) = 2.04, *p* = .042). We found that overall, patients showed a trend to vary more in their within-task accuracy than controls (LME A.3: Group *t*(100) = 1.97, *p* = .052). Accuracy in both groups varied more in the second versus first run (LME A.3: Run *t*(100) = 2.33, *p* = .022), but this effect seems to be driven by an increase in variance for controls rather than patients (LME A.3: Group x Run t(100) = −3.05, *p* = .003), but only for the easiest task conditions (LME A.3: Group x Run x Difficulty *t*(100) = 2.42, *p* = .017) as across both groups, variability of accuracy decreased in the second run for the most difficult task conditions (LME A.3: Run x Difficulty *t*(100) = −2.47, *p*= .015). We found no effects of within-task variance in reaction times (LME A.4).

### Neuroadaptive Bayesian optimisation is a feasible technique for patients

With respect to our real-time optimisation results of FPN>DMN dissociation across the task space, we found significant intra-subject reliability (i.e., spatial correlation of Bayesian predictions across the task space between the two runs of each subject) for controls (median Spearman rho ± SD: 0.91 ± 0.18, *p* < .001) and patients (0.71 ± 0.45, *p* < .001). When investigating how FPN>DMN contrast values varied for the same task within an individual (i.e., when sampled multiple times), we found no significant difference in variance between patients and controls (LME B.4: Group *t*(108) = 0.86, *p*=.39). With respect to in-scanner motion, we found that both patients and controls moved significantly more in the second run (LME B.5: Run *t*(47) = 2.34, *p* = .023) but that there was no significant difference between the two groups (LME B.5: Group *t*(47) = 1.74, *p* = .088). This indicates robustness of our obtained results and demonstrates the feasibility of the approach to achieve reliable results in patient populations.

### Semantic Judgement, Calculation and Encoding tasks maximally dissociate FPN from DMN in patients and controls

Group-level Bayesian predictions across the task space (i.e., GP regression on all observations) are shown in Fig. 3a for patients and controls, separately. We found that across both groups, Semantic Judgement (LME B.1: *t*(771) = 5.11, *p*<.001), the Calculation (LME B.1: *t*(771) = 4.39, *p* < .001) and Encoding (LME B.1: *t*(771) = 4.26, *p*<.001) tasks maximally differentiate the FPN from the DMN. Collapsed over all tasks, more difficult task conditions result in a larger FPN-DMN dissociation in both groups (LME B.1: Difficulty *t*(771) = 2.95, *p* = .003). The sampling behaviour of the Bayesian optimisation algorithm clearly confirms these results (Fig. 2d): for both patients and controls the most difficult conditions of these three tasks were most often selected by the algorithm, indicating that the algorithm identified them as optimal for maximising the dissociation between FPN and DMN. While this is very pronounced for controls, in particular for difficulty level 3 of the Calculation task (Fig. 2d – middle panel); it is worth noting that the algorithms sampled much more exhaustively across the task space for patients (Fig. 2d – left panel), potentially indicating more diversity in the optima identified among individual patients. Surprisingly, at the group level, it appears that patients do not show a qualitatively different FPN-DMN dissociation pattern across the task space compared to controls (Fig. 3a), but only seem to have a slightly diminished dissociation between these two networks for the Semantic Judgement, Calculation and Encoding tasks. To confirm that this is not an artefact of the GP regression approach, we plot the median FPN>DMN dissociation values for patients and controls across the task space (Fig. 3b – first row). We observe that the Bayesian predictions appropriately capture the underlying distribution of median FPN>DMN contrast values. These qualitative observations are also confirmed statistically: patients have a significantly lower FPN>DMN dissociation only for the Semantic Judgement task independent of difficulty level (LME B.1: Group x Semantic Judgement *t*(771) = −2.11, *p* =.035). This finding may be because the patients’ group-level results are not a good representation of individual results of patients and is in line with the algorithm’s sampling behaviour. To further understand the relative contribution of both the FPN and DMN to these results we also computed the activation values for both networks across the task space separately (Fig. 3b – second and third row). However, we found no significant difference among the groups for either the FPN (LME B.2: Group *t*(771) = 1.28, *p* = .20) or DMN (LME B.3: Group *t*(794) = 0.61, *p* = .54). Finally, we were interested in understanding the relationship between neural and behavioural measures. While higher FPN>DMN contrast values were associated with longer reaction times across both groups (LME C2: FPN>DMN *t*(643) = 2.47, *p* = .014), there was no significant difference of this effect in patients (LME C2: Group x FPN>DMN *t*(643) = −1.31, *p* = .19). Further, we did not find any association between the magnitude of FPN-DMN dissociation and accuracy across (LME C1: FPN>DMN *t*(746) = 1.59, *p* = .11) or between the two groups (LME C1: Group x FPN>DMN *t*(746) = 0.35, *p* = .73).

**Figure 3:**
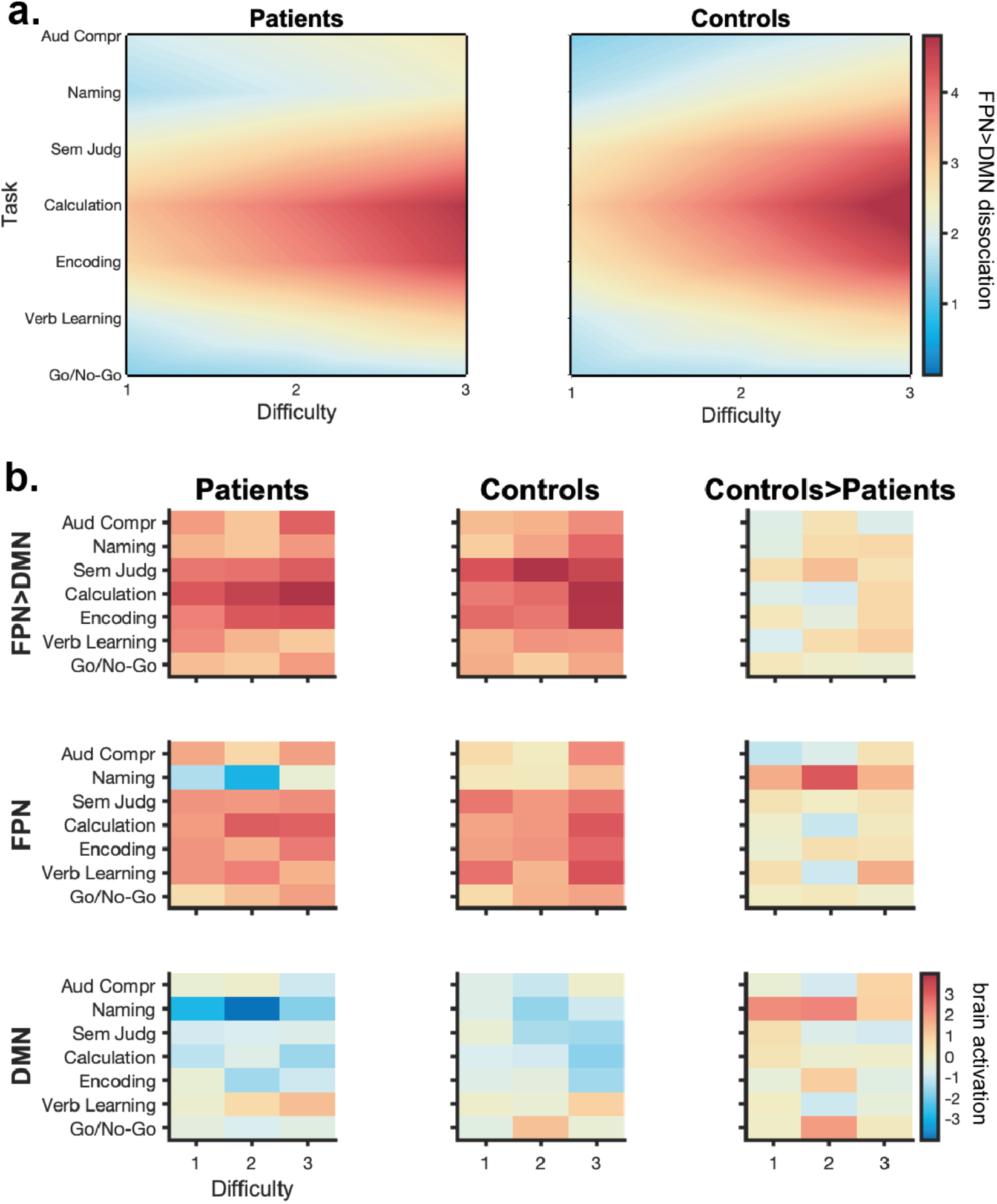
Group-level results of real-time optimisation. **(a)** Group-level Bayesian predictions across task space (i.e., GP regression across all observations) for patients (left panel) and controls (right panel) indicate no qualitative difference in the FPN>DMN dissociation pattern across the task space between both groups. Patients appear to only have a slightly diminished FPN>DMN dissociation for the Semantic Judgement, Calculation and Encoding task. **(b)** To confirm that Bayesian predictions in (a) are not driven by the specific hyper-parameters of the GP regression (see Methods), we also simply plot the median of the FPN>DMN dissociation values across the task space for both groups (first row). We observe that the Bayesian predictions appropriately capture the underlying distribution of median FPN>DMN contrast values. To understand the relative contribution of the FPN and DMN to our group-level results, we further plot the brain activation values for those networks, separately (second and third row).

### Patients show unique profiles of network dysfunction

Motivated by these findings, we wanted to understand if indeed patients’ real-time optimisation results are more diverse than controls’ results. As intra-subject reliability was high across runs, we derived each individual’s functional profile by collapsing the real-time optimisation results across both runs with the aim of deriving a more precise depiction of individuals’ functional profiles. When looking at the dissimilarity of FPN>DMN profiles (1- Spearman’s rank correlation (Kriegeskorte, 2008)) between patients, we found that they are significantly more dissimilar than the FPN>DMN profiles between controls (*t* = −5.02, *p* = .038). Interestingly, we found that patients’ individual profiles are even more dissimilar amongst each other than when comparing them with controls’ individual profiles (*t* = −2.77, *p* = .024). This indicates that patients really have unique profiles of network dysfunction but that some patients look more similar to controls than to other patients. This finding is illustrated by visualising the dissimilarity among each patient’s and control’s individual profile in 2D (Fig. 4c). We clearly see that the majority of patients group together (dark blue dots) with the exception of four particular patients (033,034, 038 and 041) who are closer to controls’ functional profiles (turquoise dots). Also, qualitatively, we observe a striking resemblance between those particular patients’ profiles and controls’ profiles (Fig. 4a and Fig. 4b), showing the highest FPN-DMN dissociation for the most difficult conditions of the Semantic Judgement, Calculation and Encoding tasks. In contrast, a single control subject (020) is grouped closer to patients than to other controls (Fig. 4c). Again, this finding is also confirmed qualitatively when inspecting this control’s functional profile and finding no resemblance with any of the other healthy controls’ functional profiles (Fig. 4b). Moreover, similar to some patients’ profiles (Fig. 4a), this control shows a high FPN-DMN dissociation for the easiest condition of the Semantic Judgement task.

**Figure 4:**
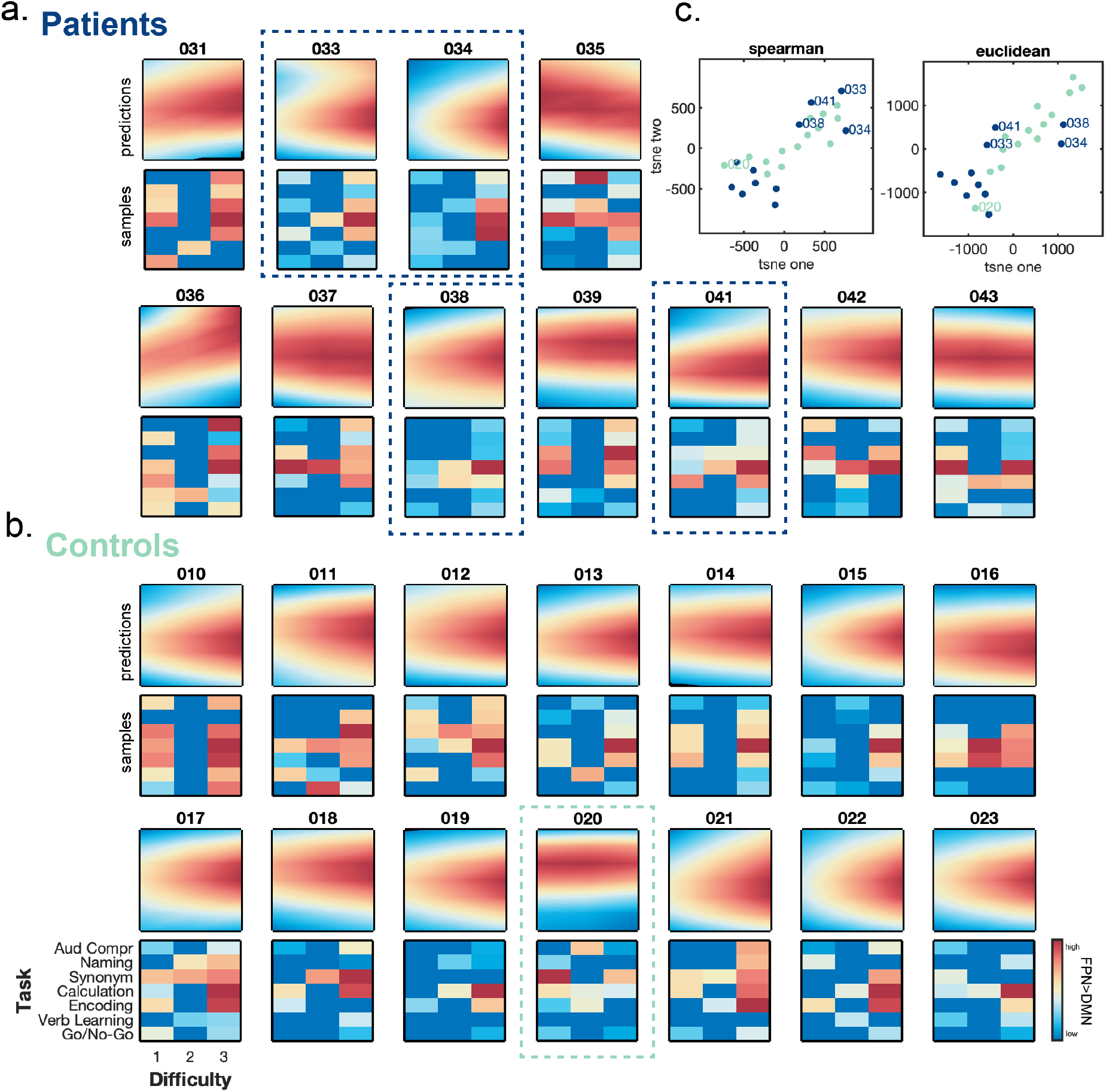
Subject-level results of real-time optimisation. **(a)** Patients show unique profiles of FPN>DMN dissociation across the task space (first row). **(b)** It can be clearly seen that in contrast to patients, controls show a striking similarity of FPN>DMN dissociation across the task space. For all patients and controls, we show the Bayesian **predictions of** FPN>DMN dissociation (i.e., GP regression based on subject-specific samples) across the entire task space as well as all **samples** individually; when a task was sampled multiple times within a subject, we computed the median across those samples. We can see that Bayesian predictions appropriately capture the underlying distribution of samples. **(c)** We used tSNE (see Methods) to visualise the dissimilarity among each patient’s and control’s individual functional profile in 2D. We found that most patients cluster together; however, four patients (i.e., 033,34,038 and 041) look more similar to controls than other patients’ functional profiles (surrounded by dotted line in dark blue). Contrarily, one control participant (i.e., 020) clusters closer with patients than other controls and indeed this control’s functional profile looks distinct from all other controls (surrounded by dotted line in turquoise).

Next, we wanted to understand if the four patients identified as more similar to controls than other patients also performed better behaviourally than all other patients. For this, we simply ranked patients by Comprehensive Aphasia Test (CAT) score, a battery of tests they performed before the experiment outside of the scanner. Results demonstrate (Table 3) that patients 033,034, 038 and 041 are ranked among the five most mildly affected patients; there is only one other patient (039) that performed equally well (ranked third) to those four patients but whose profile was not identified similar to those of controls (Fig. 4c). While these four patients (i.e., 033,034, 038 and 041) are characterised by small lesion sizes (i.e., < 5 cm^3^, Table 3), it can be seen that lesion size does not solely explain our results since patients 036 and 039 also have small lesions but show distinctively different functional profiles from these four patients.

Since our patient cohort suffers from chronic post-stroke aphasia, we would expect that the result of patients exhibiting unique patterns of network function is not specific for the dissociation of the FPN from the DMN but also holds for functional networks classically associated with language. To test this assumption, we repeated the above-mentioned analyses for a left-lateralized language network (i.e., Component 05 from (Yeo et al., 2014)). As hypothesised, we found the same results: patients’ language network profiles are more dissimilar to each other than are controls to each other (*t* = −6.04, *p* = .017). Patients are also more dissimilar among each other than when comparing them with controls’ profiles (*t* = −2.53, *p* = .024). By contrast, when focusing our analysis on a specific functional network associated with motor function (i.e., Component 01 from (Yeo et al., 2014)), we do not find that patients’ profiles compared to each other are more dissimilar than controls compared to each other (*t* = 0.1, *p*=.568), or that patients are more dissimilar to each other than when compared to controls (*t* = −0.45, *p* = .382). These supporting analyses illustrate our method’s specificity in characterising individual-level network dysfunction in patients.

### There is no single task that differentiates patients from controls

Functional profiles derived from real-time optimisation seem suitable for differentiating severely affected stroke patients from mildly affected stroke patients and healthy controls (Fig. 4c), indicating their potential usefulness as clinically relevant biomarkers. To further understand if these multivariate profiles of residual network function yield information we could have not obtained from univariate analyses of individual tasks as conventionally performed in clinical neuroimaging research, we “normalised” each patient’s task-specific Bayesian prediction values with respect to the controls’ distribution of FPN>DMN Bayesian predictions for that particular task. Results from this analysis reveal on which particular task each patient significantly deviates from healthy controls. Importantly, we found that no single task exists for which all patients significantly differ from controls, or even where the direction of difference is the same across patients (Fig. 5a). While many patients show a diminished FPN>DMN dissociation for medium and difficult conditions of the Semantic Judgment, Calculation, or Encoding task, this effect is only significant in two patients (035,036). Surprisingly, other patients show a reverse pattern and a significant stronger FPN>DMN dissociation for exactly the same tasks (034, 038). Different to our group-level results, the most difficult level of the Verbal Learning task appears to be the task for which the majority of our patients (i.e., six) show a significantly different FPN>DMN contrast value; however, for only four (031, 035, 036, 042) of these patients the FPN>DMN dissociation is diminished compared to controls; for the other two patients (034, 038), we observed the reverse and they showed a stronger FPN>DMN dissociation than controls. For comparison, we repeated this analysis for each controls’ individual functional profiles (i.e., normalised each to their own group-level results); results of these analyses are shown in Fig. 5b. We found that most controls show either negligible differences or a stronger FPN>DMN dissociation for medium and difficult conditions of the Semantic Judgment, Calculation, or Encoding task; for three controls the stronger FPN>DMN dissociation on those tasks even reaches statistical significance (012, 016, 018). Two other controls are characterised by a significantly stronger FPN>DMN dissociation on the Go/No-Go task. Interestingly, out of 15 controls, only a single subject (020) showed a significantly diminished FPN>DMN dissociation across a wide range of tasks conditions (i.e., medium and most difficult levels of the Semantic Judgment, Calculation, Encoding task and Verbal Learning tasks). This is the same control subject identified as resembling the functional profile more similar to patients than other controls (Fig. 4c).

**Figure 5:**
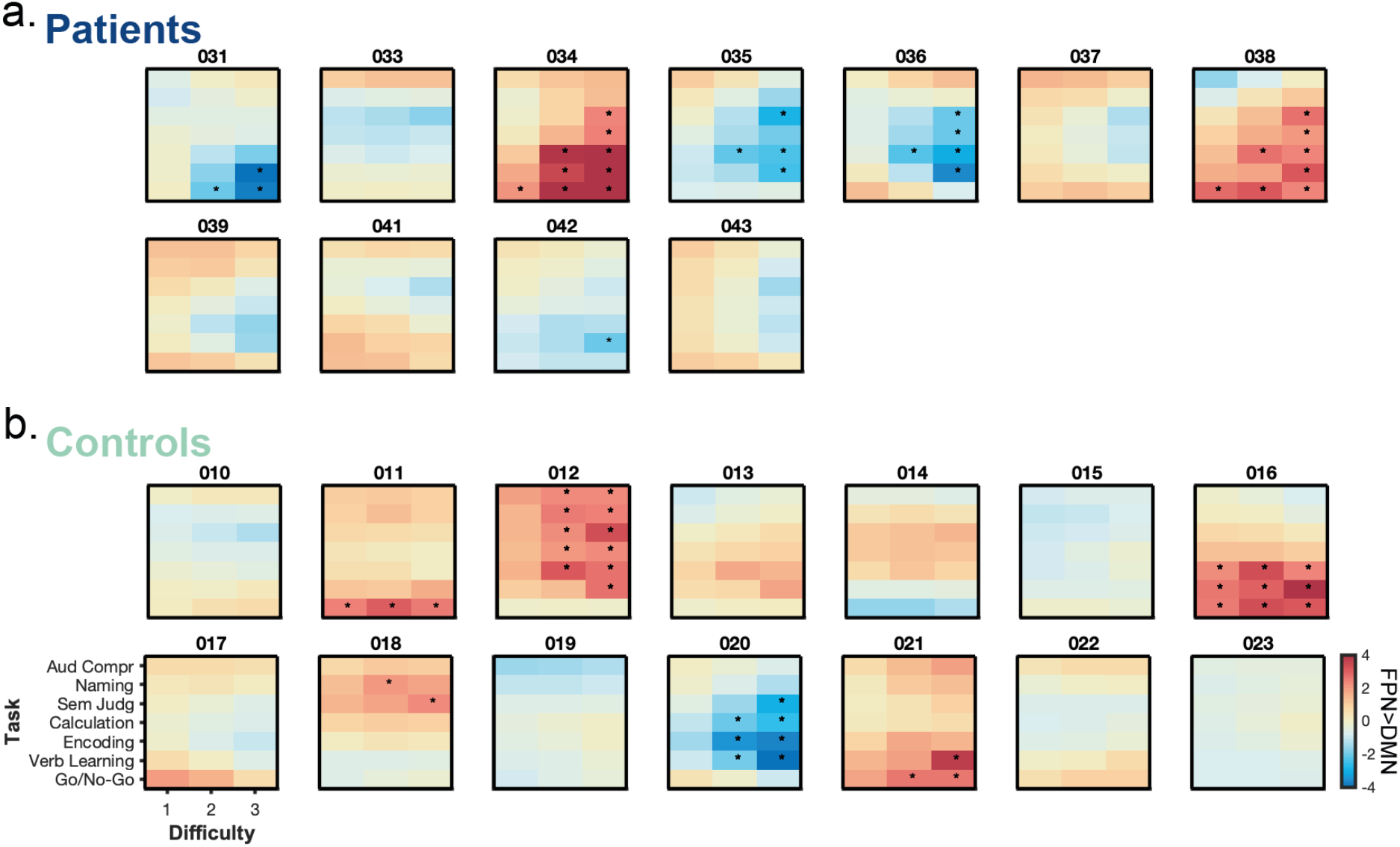
Individual functional profiles normalised to controls’ group result. **(a)** When looking at patients’ normalised functional profiles, we found that no single task exists, for which all patients significantly differ from controls or even where the direction of difference is the same for all patients. It is even the case that for a few tasks, some patients show a significantly diminished (i.e., 35, 36, 31) FPN>DMN dissociation while the same tasks are associated with a significantly stronger FPN>DMN dissociation for other patients (034,038). **(b)** When looking at control’s normalised functional profiles, we found that they either have negligible smaller or stronger FPN>DMN dissociation compared to their group result. Only one control subject (020) shows a significantly diminished FPN>DMN dissociation for many different medium to difficult task conditions; interestingly, this is the same control subject whose functional profile was associated more with patients’ functional profiles than with other controls’ profiles (Fig. 4c).

## Discussion

In this study, we applied neuroadaptive Bayesian optimisation for the first time to a cohort of patients with the aim of rapidly searching through a variety of different cognitive and language-related task conditions in order to obtain patient-specific functional profiles of residual domain-general network function.

At the group level, patients did not show an altered FPN>DMN dissociation pattern across the task space compared to controls. For both patients and controls, more difficult task conditions and particularly the Semantic Judgement, Calculation and Encoding tasks maximally dissociated the FPN from the DMN. This is in line with our previous work, showing that the Calculation and Encoding tasks as well as increased task demands strongly recruit this particular FPN in healthy volunteers (Lorenz et al., 2018; Yeo et al., 2014). While we found no significant difference between patients and controls for the Calculation and Encoding tasks, we did find the Semantic Judgement task to be associated with a significantly diminished FPN>DMN dissociation in patients on a group level. However, when investigating individual patients’ neural responses for the Semantic Judgement task, only two out of all patients showed a significantly lower FPN>DMN dissociation compared to controls and two other patients showed a significantly stronger FPN>DMN dissociation on that task compared to controls. This highlights the limitations involved with the conventional approach in clinical neuroimaging: it is remarkably difficult to predict before the start of a clinical study which task will reveal a sensitive biomarker that can be applied to an individual patient and that is capable of differentiating patients from controls as well as subgroups of patients; this is especially important considering that patients may have multiple co-morbidities (e.g., vascular disease) that differentially affect brain networks.

Our findings clearly demonstrate that group-level results are not representative of individual patient-specific results. We found greater heterogeneity among the functional profiles of patients than among those of controls. In fact, patients feature idiosyncratic profiles of FPN>DMN dissociation across the task space. This was also in line with the sampling trajectories of the real-time algorithm that sampled much more diversely for patients. By contrast, the algorithm’s sampling was much more focused for controls as they showed a very high consistency in their functional profiles; a finding replicating our earlier work (Lorenz, Monti, Violante, et al., 2016; Lorenz et al., 2018). Interestingly, we found that heterogeneity among the functional profiles of patients was even higher than when looking at the heterogeneity between patients and controls. We could show that, indeed, four of our patients had functional profiles more closely resembling those of age-matched healthy controls than those of other stroke patients. Importantly, these four patients were among the top five performing patients in an exhaustive out-of-scanner test battery. We confirmed the validity of patient-specific functional profiles by comparing the real-time optimisation results of two independent runs. We found patient-specific profiles to be consistent; however, controls’ functional profiles were characterised by a much higher intra-subject reliability. This lower intra-subject reliability of patient-specific functional profiles may be explained by patients showing learning effects (improved accuracy) in the second run, potentially contributing to slightly different results across both runs.

One challenge of this closed-loop experimental framework is that subjects are required to remember various task instructions and switch between tasks in a relatively swift manner (every minute in our case). Despite these heightened task demands, in our study, patients performed above chance for all tasks on at least the easiest difficulty level (an exception was the Verbal Learning task for which also controls performed at chance, see Methods). For some tasks, patients failed to perform above chance for the most difficult task conditions (Semantic Judgement, Calculation and Encoding tasks); in the future, it should be considered to lower the difficulty (or consider a wider range of difficulties) for those tasks to make them feasible for patients. In addition, task instruction at each iteration could be provided both orally and visually as well as presented for longer as some patients encountered problems with the length of presentation of instruction slides. While patients varied more in their behavioural performance on the same tasks than controls, this effect was not reflected in the neural measures. Interestingly, patients’ accuracy increased in the second run, illustrating that patients get acquainted with the experimental paradigm and that they could potentially benefit from an in-scanner practice run before the real-time experiment. On the other hand, patients (and controls) moved more in their second run, highlighting the need for carefully trading-off longer practice time in the scanner and reducing motion artefacts to improve data quality.

In summary, our study highlights the importance of moving beyond traditional “one-size-fits-all” approaches in clinical neuroimaging where patients are treated as one group and single tasks (or a few tasks) are used. Instead, we demonstrate that mapping residual network activity following brain injury across many different tasks using real-time optimisation yields robust patient-specific functional profiles that carry meaningful information about a patient’s behavioural capacity. From a conceptual point of view, this multi-task approach also improves the generalizability of our findings (Yarkoni, 2019). The Bayesian model that is iteratively built up, explicitly models relationships between cognitive tasks; the results from the GP regression model are, therefore, not specific to a single task but inherently estimate performance across all tasks, allowing for far more principled inferences about generalisation and the relationship between the dysfunction of brain networks and underlying cognitive processes. Thus, multivariate functional profiles of residual brain function derived from neuroadaptive Bayesian optimisation may have promising potential to become clinically relevant and generalizable biomarkers with satisfactory test-retest reliability.

A detailed understanding of how patient-specific functional profiles relate to different aspects of behavioural functioning is beyond the scope of the present study and needs to be addressed in studies with larger sample size and an even wider variety of cognitive tasks and conditions. As we found no univariate linear relationship between FPN>DMN dissociation and task accuracy in this study, we argue that the relationship between patient’s profiles of network function and behaviour is most likely of multivariate and possibly also non-linear nature. In the future, it would be important to test patients who undergo real-time optimisation also on a large battery of different tasks outside of the scanner. Multivariate patient-specific functional network profiles could then be related to multivariate behavioural profiles (e.g., (Butler et al., 2014; Halai et al., 2017)) using, e.g., canonical correlation analyses or other machine learning techniques.

In addition, longitudinal studies need to be carried out in order to advance our understanding whether and how functional profiles derived from neuroadaptive Bayesian optimisation predict stroke recovery and thus, could be used to guide rehabilitation strategy: so far it is not clear if therapy should focus on tasks associated with a high residual FPN>DMN dissociation, low residual FPN>DMN dissociation or tasks for which residual FPN>DMN dissociation is most different to controls. Interestingly, our approach can be combined with therapeutic interventions involving non-invasive brain stimulation (Lorenz et al., 2019; Lorenz, Monti, Hampshire, et al., 2016). Using neuroadaptive Bayesian optimisation, cognitive task conditions and non-invasive brain stimulation parameters could be searched through simultaneously (e.g., designing a two-dimensional search space with type of task along one dimension and stimulation intensity along the second) with the aim of identifying optimal therapeutic protocols tailored to individual patients for behavioural therapy (i.e., optimal task) in conjunction with brain stimulation (i.e., optimal stimulation intensity).

Another avenue for future research would be to incorporate lesion information in a systematic manner. Structural brain imaging has been shown to predict stroke patients’ current linguistic and cognitive impairments (Halai et al., 2020) and language outcome and recovery (Hope et al., 2013; Seghier et al., 2016). One avenue for future work would be to combine measures derived from structural scans (~ 5 min) with rapidly obtainable functional profiles (~ 10-15 min) to further boost the accuracy of such predictions.

The strength of our real-time optimisation approach lies in the rapid mapping out of functional profiles of residual network function across a large space of cognition without the need to exhaustively sample all possible tasks. This efficiency makes it a highly interesting tool for clinical populations; yet it may come at a cost of sensitivity. For example, we identified four stroke patients whose functional profiles are similar to those of healthy controls. Given our relatively coarse sampling across the task space, it may be possible that we were not able to pick up fine-grained differences between the functional profiles of high-functioning stroke patients and age-matched healthy controls. Therefore, an interesting future direction may be to use neuroadaptive Bayesian optimisation as a first stage for obtaining a comprehensive yet coarse depiction of residual brain function.

Results obtained from this first stage could then be used to inform a second stage of dense sampling (DiNicola et al., 2020; Gordon et al., 2017; Poldrack et al., 2015). Patients could then be tested repeatedly over a long period of time on a subset of tasks identified with real-time optimisation, or real-time optimisation repeated repeatedly (e.g., (Lorenz et al., 2018)). Such a two-stage procedure would yield very precise individual functional profiles across the most informative tasks, and allow a better grasp on whether differences observed reflect trait or state alterations in functional brain networks and potential for therapeutic interventions.

From a methodological point of view, robust online stopping criteria for the real-time optimisation algorithm need to be developed. In this study, the number of task block iterations per run was predetermined; however, one avenue for future work is to automatically end the run only when the uncertainty of the algorithm over the task space is sufficiently small and/or enough statistical evidence has been accumulated. This could result in more accurate functional profiles (the algorithm samples more in face of uncertainty) as well as greater efficiency of the approach (the run could be stopped earlier). While we have proposed two online stopping criteria in the past that rely on characteristics of the acquisition function (Lorenz et al., 2015), more work is needed to assess how well these perform in clinical populations.

In conclusion, we show for the first time, that neuroadaptive Bayesian optimisation is a feasible, reliable and highly efficient approach for identifying patient-specific functional profiles of network dysfunction across many different task conditions. While the sample size is currently small, we show that these unique patient profiles are associated with behaviour; thereby demonstrating the potential of this approach for exploring and testing novel neuroimaging biomarkers for recovery after stroke. Importantly, this approach can be extended to optimise for any target brain network/state (e.g., functional connectivity or multivariate pattern) and optimise task conditions and non-invasive brain stimulation parameters conjointly, thereby, opening new avenues for precision medicine for a wide range of different neurological and psychiatric conditions.

## Methods

### Participants

The study was approved by the National Research Ethics Service Committee. We recruited 14 patients with left hemisphere infarcts, over the age of 40 and premorbid fluency in English. Patients with a previous history of a stroke resulting in aphasia or other neurological illness, or concurrent use of psychoactive drugs, were excluded. The average lesion volume was 10.56 cm^3^ ± 8.03 cm^3^. Table 2 contains further patient details. As controls, we recruited 15 fluent English speaking, healthy participants, over the age of 40. For the pilot study, 8 younger and fluent English speaking, healthy participants were recruited. Controls and pilot participants had no history of any neurological/psychiatric disorders. All study participants (patients, controls and pilot participants) were right-handed, had normal or corrected-to-normal vision and normal adult hearing. Participants were informed about the real-time nature of the fMRI experiment, but no information was given on the actual aim of the study or which parameters would be adapted in real-time. The investigator was not blinded due to the complexity of data acquisition and the need to ensure that real-time optimisation was functioning.

**Table 2:**
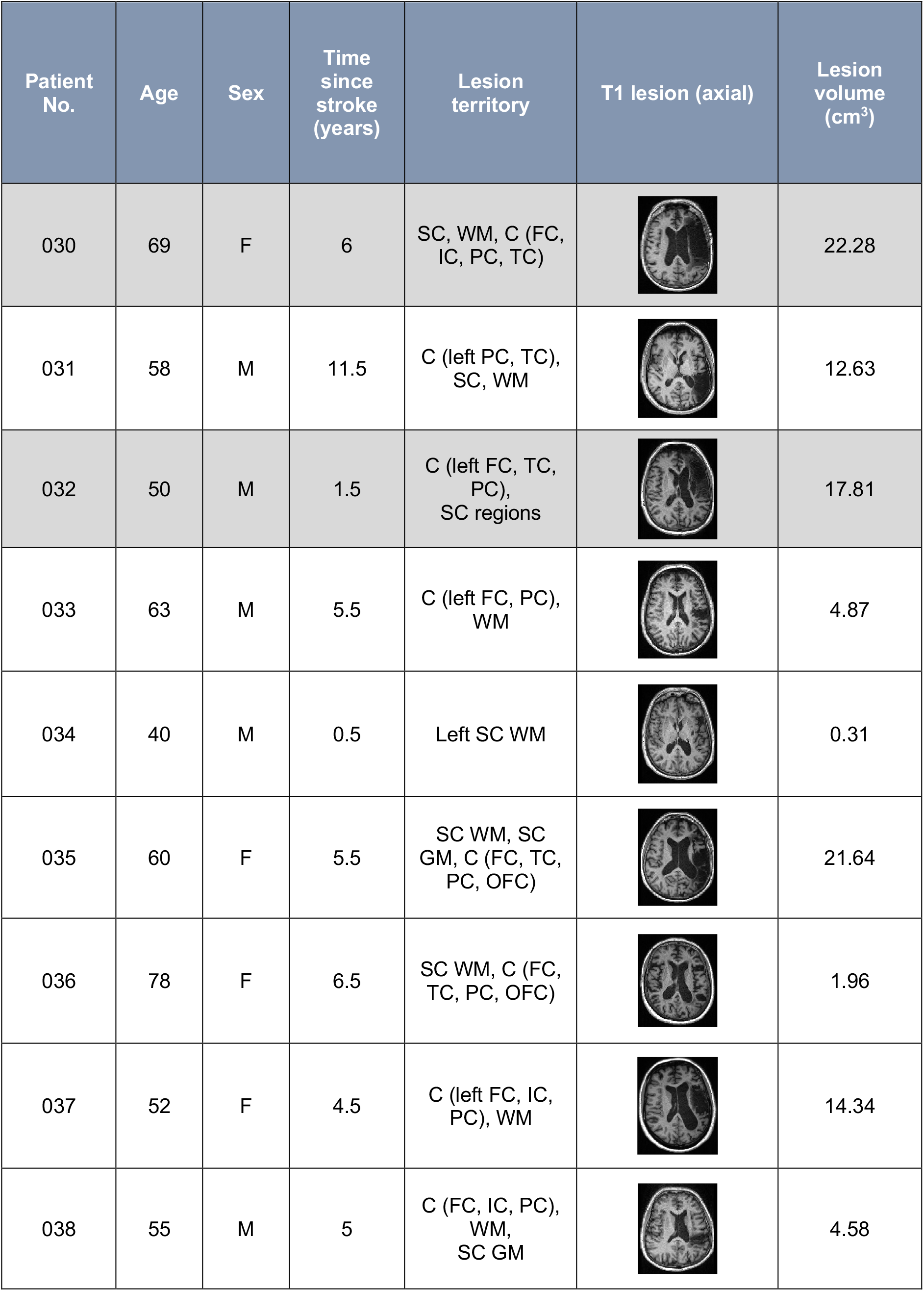
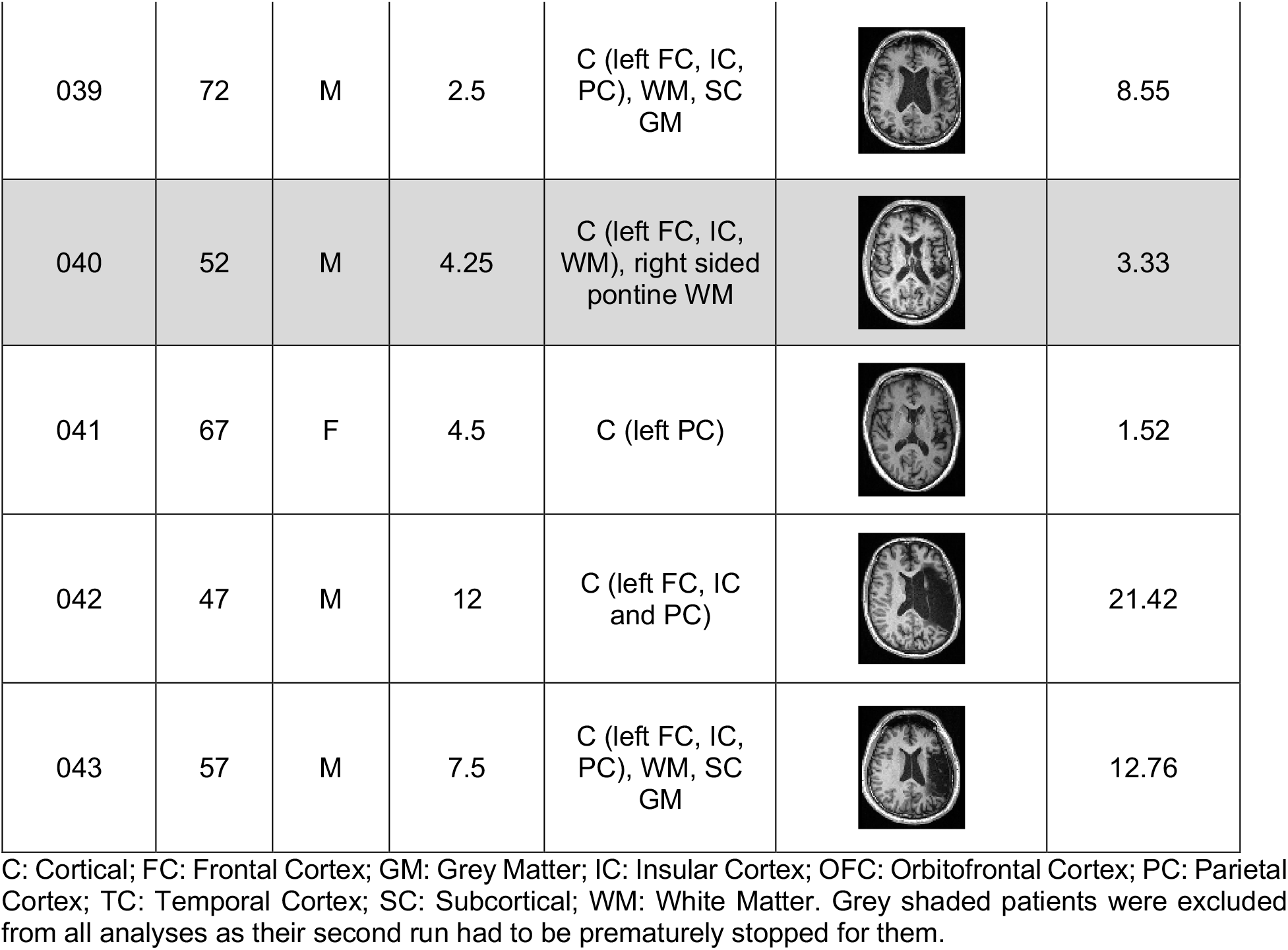
Details of stroke patients.

| Patient No. | Age | Sex | Time since stroke (years) | Lesion territory | T1 lesion (axial) | Lesion volume (cm<sup>3</sup>) |
| --- | --- | --- | --- | --- | --- | --- |
| 030 | 69 | F | 6 | SC, WM, C (FC, IC, PC, TC) |  | 22.28 |
| 031 | 58 | M | 11.5 | C (left PC, TC), SC, WM |  | 12.63 |
| 032 | 50 | M | 1.5 | C (left FC, TC, PC), SC regions |  | 17.81 |
| 033 | 63 | M | 5.5 | C (left FC, PC), WM |  | 4.87 |
| 034 | 40 | M | 0.5 | Left SC WM |  | 0.31 |
| 035 | 60 | F | 5.5 | SC WM, SC GM, C (FC, TC, PC, OFC) |  | 21.64 |
| 036 | 78 | F | 6.5 | SC WM, C (FC, TC, PC, OFC) |  | 1.96 |
| 037 | 52 | F | 4.5 | C (left FC, IC, PC), WM |  | 14.34 |
| 038 | 55 | M | 5 | C (FC, IC, PC), WM, SC GM |  | 4.58 |
| 039 | 72 | M | 2.5 | C (left FC, IC, PC), WM, SC GM |  | 8.55 |
| 040 | 52 | M | 4.25 | C (left FC, IC, WM), right sided pontine WM |  | 3.33 |
| 041 | 67 | F | 4.5 | C (left PC) |  | 1.52 |
| 042 | 47 | M | 12 | C (left FC, IC and PC) |  | 21.42 |
| 043 | 57 | M | 7.5 | C (left FC, IC, PC), WM, SC GM |  | 12.76 |

C: Cortical; FC: Frontal Cortex; GM: Grey Matter; IC: Insular Cortex; OFC: Orbitofrontal Cortex; PC: Parietal Cortex; TC: Temporal Cortex; SC: Subcortical; WM: White Matter. Grey shaded patients were excluded from all analyses as their second run had to be prematurely stopped for them.

**Table 3:** Patient’s ranking based on Comprehensive Aphasia Test (CAT) scores and their lesion volume (a higher score corresponds to better language ability)

| Patient | Ranked CAT score (sum) | Lesion volume (cm<sup>3</sup>) |
| --- | --- | --- |
| 034 | 805.0 | 0.31 |
| 038 | 702.5 | 4.58 |
| 039 | 627.5 | 8.55 |
| 033 | 595.0 | 4.87 |
| 041 | 590.5 | 1.52 |
| 036 | 573.5 | 1.96 |
| 037 | 558.0 | 14.34 |
| 031 | 511.0 | 12.63 |
| 035 | 424.5 | 21.64 |
| 042 | 422.5 | 21.42 |
| 043 | 179.0 | 12.76 |

### Out-of-scanner behavioural assessment

All patients underwent the Comprehensive Aphasia Test (CAT) outside of the scanner (Swinburn et al., 2005) before the experiment started. This provided a measure for word fluency, comprehension of spoken language, comprehension of written language, repetition, naming, reading, writing, descriptive speaking, descriptive writing, as well as a cognitive score. All the scores were summed to provide a single measure of aphasic deficit for the patients.

### Task space

A two-dimensional task space was designed consisting of seven different tasks with three difficulty levels each. All tasks and their variants are briefly described below and depicted in Fig. 6.

**Figure 6:**
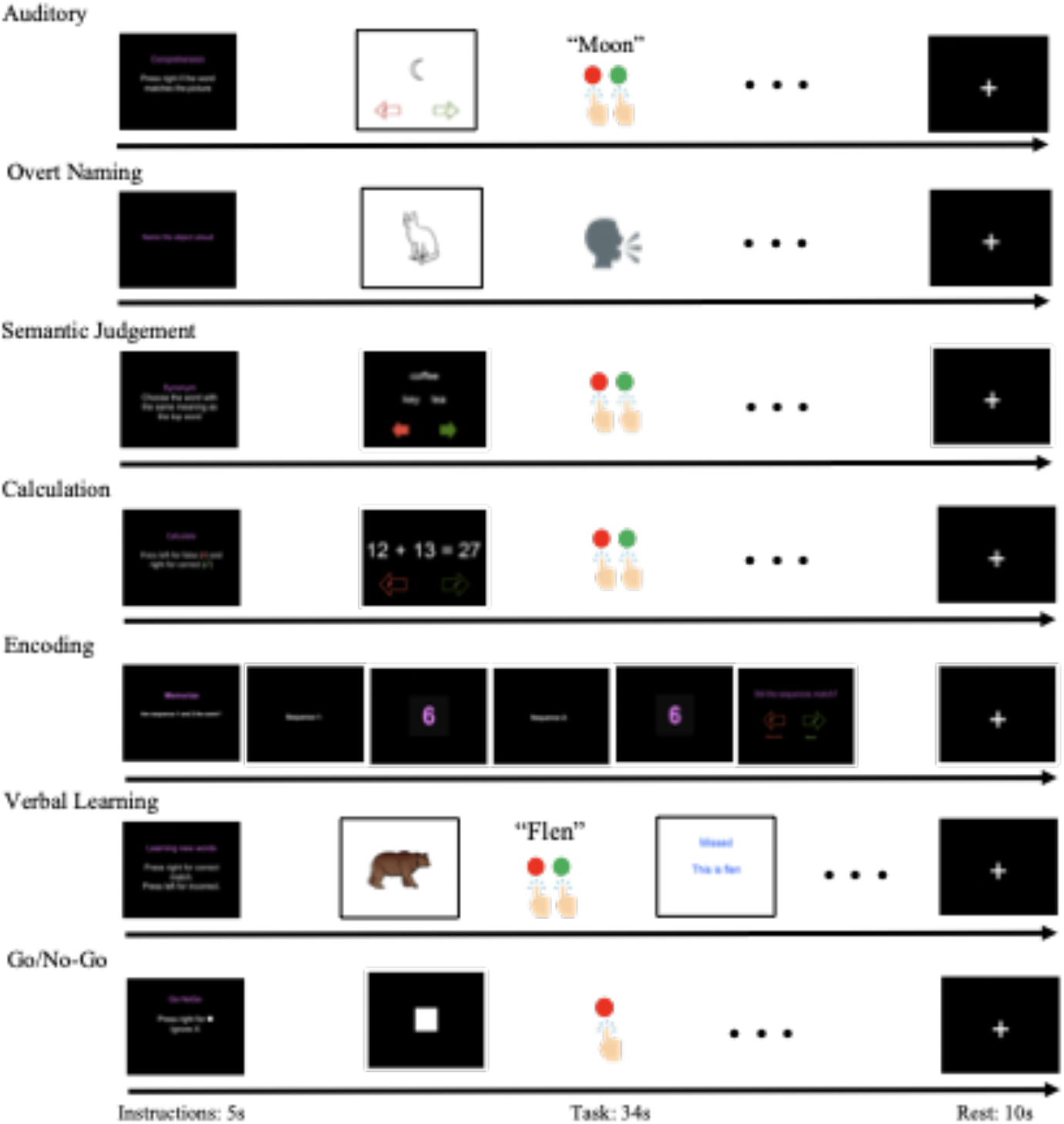
Stimulus presentation of each task. Each task block lasted 34 s followed by 10 s rest. Preceding each task block, participants received a brief instruction (5 s) about the task they would need to perform in the upcoming block followed by a short 3 s rest period (black background, not shown here). Except the Overt Naming tasks, all tasks required patients pressing buttons to indicate their response.

#### Auditory Comprehension

This language task tested participants’ ability to understand verbal word stimuli and match it to the correct picture. Correct pairings were indicated using a keypad. The word picture associations were based on the PALPA (Psycholinguistic Assessments of Language Processing in Aphasia). At difficulty level 1, 10 image-spoken word pairs were presented and pairings were always correct. At difficulty level 2, 14 image-spoken word pairs were presented and unrelated distractors were included. The probability of receiving an incorrect pairing was 50%. At difficulty level 3, close semantic distractors were included and the probability of receiving an incorrect pairing was 50%. Each image was shown for 3 s.

#### Overt Naming

This language task tested participants’ ability to name objects. Participants were shown a series of images and required to verbalise (into a microphone) what the stimulus represented. Responses were listened to online to ensure participants performed the task as instructed, by attempting to verbalise at the correct times. Level 1 included easy words and level 2 included hard words. Stimuli were classified as easy or hard based on reaction times and name agreement data collected in a previous experiment (Krishnan et al., 2015). At level 3, 25 words with the lowest frequency were used from the Graded Naming Test (McKenna & Warrington, 1983). As this task did not require any button responses and rather relied on the quality of the verbal output, chance level could not be determined.

#### Semantic Judgement

This language task tested participants’ ability to decipher the meaning of words. It was designed by (Jefferies et al., 2009) and included words scaled by their interpretation difficulty, based on imageability and frequency. Participants were required to indicate which two words (out of three) were semantically related using a keypad. Each trial appeared on screen for 3.5 s. At higher difficulty levels the words chosen had lower frequency and imageability.

#### Calculation

This non-language task tested participants’ ability to perform mental arithmetic. Patients with aphasia often suffer from acalculia (Luccia & Ortiz, 2016), and this task is also known to rely heavily on the FPN (Lorenz et al., 2018; Yeo et al., 2014). In each equation, participants had to decide whether the answer provided was correct or incorrect using the keypad. At level 1, the equations involved one-digit addition and each equation was shown for 2.9 s, at level 2, it involved addition of numbers between 10–20. At level 3, it involved higher two-digit subtraction and each equation was shown for 3.5 s. The probability of receiving an incorrect answer was 50%.

#### Encoding

This non-language task tested participants’ working memory through digit span. Each digit in a sequence appeared on a black background for 0.4 s with the next digit in the sequence appearing after a 0.1 s gap. This was then followed by a second sequence of numbers of the same length. Participants were required to indicate whether the second sequence matched the first using a keypad. At level 1, digit span was three, at level 2, digit span was 5 and at level 3, digit span was seven. The probability of receiving a mismatched number sequence was 50%.

#### Verbal Learning

This language task tested participants’ ability to learn new words based on feedback. It was designed by (Sliwinska et al., 2017) and was shown to rely heavily on the FPN. Participants were presented with a series of images with audio stimuli denoting a pseudo word (using English phonology) to each image. Without prior training, participants were required to indicate whether the image and word pairing was correct. They were then provided with feedback as to whether their response was correct and provided with the correct name. If they did not respond, they still received feedback (Sliwinska et al., 2017). Level 1 involved learning two correct pseudo word–object associations, level 2 involved learning four and level three involved learning 10. At each level 3, correct associations were mixed with 25 incorrect associations. However, with hindsight this task was not suited for the current study as for many pairings (in particular for level two and three), participants were not able to correctly learn the new pairings within the short blocks of 34 s. As a result, patients and healthy controls performed at chance on this task.

#### Go/No-Go

This non-language task tested participants’ ability to identify targets and inhibit responses. In each trial either a white square, which required a “Go” response (pressing the right key on the keypad), or a cross which required a “No-Go” response (participants did nothing) were shown on screen. At level 1, stimuli were presented for 0.5 s with a 1 s gap, and 20% of trials were “Go”. At level 2, stimuli were presented for 0.25 s with a 0.75 s gap, and 35% of trials were “Go”. At level 3, a blue square was also incorporated, which participants were supposed to treat as a “No-Go” as well as the white cross. Stimuli were presented for 0.25 s with a 0.7 second gap, and 30% of trials were “Go”.

### Experimental procedure

Each participant underwent two separate real–time optimisation runs. The task space and optimisation target (i.e., FPN>DMN) were identical across the two runs. Each run was initiated randomly (i.e., first four task blocks were selected randomly from across the task space) and no data from previous runs/subjects were entered into the algorithm. The start of each run was synced with the onset of the first TR and each new task block was initiated by a TR. The first task commenced after 10 TRs to allow for T1 equilibration effects. Each run lasted 14.2 min and consisted of 16 task block iterations; each iteration consisted of a task block lasting 34 s followed by 10 s rest block (white fixation cross on black background). Preceding each task block, participants received a brief instruction (5 s) about the task they would need to perform in the upcoming block followed by a short 3 s rest period (black background). For five patients, task instructions had to be provided orally via a microphone due to reading impairments. Where participants were required to use a keypad to indicate answers, they used their left hand to avoid difficulties related to right sided motor impairments (as is common in left middle cerebral artery (LMCA) stroke). The keypad had two buttons, the right button for correct answers and the left button for incorrect answers. Where only one button response was required the right was pressed (Fig. 6). Auditory stimuli were presented using sound-attenuating in-ear MR-compatible headphones (Sensimetrics, Model S14, Malden, USA). Subjects were trained and familiarised with all tasks outside of the scanner prior to the start of the experiment. For three patients (030, 032 and 040) the second run was discarded due to distress and/or fatigue, causing us to stop the run prematurely. One control (009) failed to hear the auditory stimuli; all other participants indicated that they could hear the words during a test run in the scanner.

### FPN and DMN network masks

Masks of the bilateral target brain networks were based on a meta-analysis reported in (Yeo et al., 2014). For the FPN, we selected Component 09 covering the superior parietal cortex, the intraparietal sulcus, the lateral prefrontal cortex, anterior insula and temporoparietal junction. For the DMN, we selected Component 11, spanning the anterior hippocampus, posterior cingulate cortex and medial prefrontal cortex. Thresholded (z > 2) and binarized maps of the two brain networks were used as masks for our real-time study.

### Real-time fMRI

For real-time fMRI data processing, we followed a similar procedure as described in our previous work (Lorenz et al., 2018). Images with whole-brain coverage were acquired in real-time by a Siemens Verio 3 T scanner using an EPI sequence (T2*-weighted gradient echo, voxel size: 3.00 × 3.00 × 3.00 mm, field of view: 192 × 192 × 105 mm, flip angle: 80°, repetition time (TR)/echo time (TE): 2000/30 ms, 35 interleaved slices). Prior to the online run, a high-resolution gradient-echo T1-weighted structural anatomical volume (voxel size: 1.00 × 1.00 × 1.00 mm, flip angle: 9°, TR/TE: 2300/2.98 ms, 160 ascending slices, inversion time: 900 ms) and one EPI volume were acquired. Online pre-processing was carried out with FSL (Jenkinson et al., 2012). The first steps occurred offline prior to the real-time fMRI scan. Those comprised brain extraction using BET (Smith, 2002) of the structural image followed by a rigid-body registration of the functional to the downsampled structural image (2 mm) using boundary-based registration (Greve & Fischl, 2009) and subsequent affine registration to standard brain atlas (MNI) (Jenkinson et al., 2012; Jenkinson & Smith, 2001). In patients with large lesions, BET was performed iteratively with various options (e.g., centre-of-gravity, fractional intensity threshold, vertical gradient in fractional intensity threshold) to allow the best brain extraction; results were inspected visually by experimenter. The resulting transformation matrix was used to register the FPN and DMN from MNI to the functional space of the respective subject. For online runs, incoming EPI images were motion corrected (Jenkinson et al., 2002) in real-time with the previously obtained functional image acting as reference. In addition, images were spatially smoothed using a 5 mm FWHM Gaussian kernel. For each TR, means of the two brain networks were extracted. To remove outliers, scrubbing (i.e., data replacement by interpolation) was performed on these time courses with the cut-off set to ± 4 SD. Removal of low-frequency linear drift was achieved by adding a linear trend predictor to the general linear model (GLM). To further correct for motion, confound regressors were added to the GLM consisting of six head motion parameters and a binary regressor flagging motion spikes (defined as TRs for which the framewise displacement exceeded 1.5).

### FPN-DMN dissociation target measure

After presentation of each task block, we calculated the difference in brain activation between the FPN and DMN. For this purpose, we ran incremental GLMs on the pre-processed time courses of each network separately. In our case, incremental GLM refers to the design matrix growing with each new task block, i.e., the number of timepoints as well as the number of regressors increasing with the progression of the real-time experiment. The GLM consisted of task regressors of interest (one regressor for each task block), task regressors of no interest (e.g., 5 s instruction period) as well as confound regressors as described above (seven motion and linear trend regressor) and an intercept term. Task regressors were modelled by convolving a boxcar kernel with a canonical double-gamma hemodynamic response function (HRF). After each new block, the beta coefficients were re-estimated. We computed the difference between the estimates of all task regressors of interest (i.e., beta coefficients) for the FPN and DMN (i.e., FPN > DMN). The resulting contrast values were then entered into the Bayesian optimisation algorithm. An initial burn-in phase of four randomly selected tasks was employed, i.e., the first GLM was only computed at the end of the fourth block after which the closed-loop experiment commenced.

### Bayesian optimisation

Bayesian optimisation is a two-stage procedure that repeats iteratively in a closed loop. The first stage is the *data modeling stage*, in which the algorithm uses all available samples obtained from real-time fMRI (i.e., FPN>DMN contrast values from the GLM) up to that iteration to predict the subject’s brain response across the entire task space using Gaussian process (GP) regression (Brochu et al., 2010; Rasmussen & Williams, 2006; Shahriari et al., 2016). GPs are fully specified by their mean and covariance functions. We employed a zero mean function and as covariance function we chose the squared exponential kernel (Rasmussen & Williams, 2006):

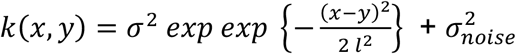

where *x*, *y* ∈ *R*^2^ correspond to the choice of task condition. The hyper-parameters *σ* ∈ *R* and *l* ∈ *R*^2^ each determine the variance and length scale of the covariance kernel, respectively. Further, it is assumed that observations are corrupted by white noise, 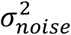. These parameters need to be selected prior to running the experiments. We used independent data from eight pilot subjects to tune these parameters using Type-2 maximum likelihood (Rasmussen & Williams, 2006). This choice of hyper-parameters was then fixed for all participants’ runs in the real-time study and the same hyper-parameters were also used for all analyses reported here. The first four task blocks served as a burn-in for a first estimate of the GP.

The second stage is the *guided search stage*, in which an acquisition function is used to propose a point in the task space to sample next (i.e., the task, the subject will need to perform in the next iteration). There are different acquisition functions available and the choice of acquisition function encodes the bias towards exploration (i.e., sampling tasks with high uncertainty) or exploitation (i.e., sampling tasks that are predicted to be optimal) (Lorenz et al., 2017). Here we used the upper-confidence bound (GP-UCB) acquisition function (Srinivas et al., 2010):

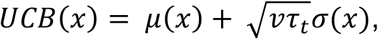

where

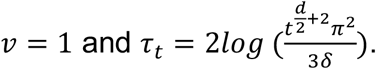

Simply put, the GP-UCB favours the selection of points with high predicted mean value (i.e., optimal tasks), but equally prefers points with high variance (i.e., tasks worth exploring). Formally, at every iteration, the next task is selected by maximising the GP-UCB:

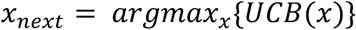

This new sample will then be used to return to the first stage and update the GP; thereby closing the experiment loop.

### Behavioural accuracy

To assess if patients understood task instructions and performed higher than chance-level on the various tasks, we computed the nonparametric effect size between the true empirical distribution of patients’ accuracy and the generated chance-level distribution for each task condition separately. The chance-level distribution was derived by randomly shuffling (1000 permutations) the trial sequence and the corresponding behavioural responses of each block, and then re-computing block-wise accuracy. This approach has the advantage of preserving the overall response pattern of each patient. For nonparametric effect size computation, we chose the area under the receiver operating characteristic curve (AUROC) approach. AUROC can be understood as a measure of overlap between two distributions and its values range from 0 to 1. A value at 0.5 indicates that there is no effect found between the two distributions. Significance was determined when the one-sided lower 95% confidence bound was higher than an AUROC of 0.5, indicating an effect between our empirically obtained and generated null distribution. Confidence intervals were computed via bootstrapping (10,000 replications). AUROC computations and bootstrapping were performed using the “Measures of Effect Size” Matlab toolbox (Hentschke & Stüttgen, 2011).

### Linear mixed effect models of behavioural and fMRI data

To assess behavioural performance, fMRI measures, in-scanner motion and the relationship between fMRI and behaviour, several linear mixed-effect (LME) models were performed. As difficulty level 2 was sampled much fewer times than difficulty level 1 and 3 (Fig. 2d), results from difficulty levels 1 and 2 were merged for this analysis. In the different models, categorical regressors were “Group” (patients or controls), “Subject” (our 25 different subjects), “Run” (run 1 or run 2) and “Task” (the seven tasks included in our task space). Ordinal regressors were “Difficulty” (difficulty level 1 or 3), and “Repetition” (corresponding to the number of times the same task has been sampled before in an individual subject’s run). Dependent variables were: accuracy (i.e., percentage of correct trials within a task block), reaction times (only of correct responses), in-scanner motion (mean of framewise displacement (FD) per run), FPN>DMN contrast values, as well as the FPN and DMN brain activation values, separately. In addition, to understand variability of behavioural and neural measures for repetitions of a specific task within an individual (i.e., when the algorithm selected the same task to be sampled multiple times in a run), we computed the median absolute deviation (MAD) of accuracy, reaction times and FPN>DMN contrast values. MAD is a more robust measure of variability, more resilient to outliers in a data set than standard deviation.

For each research question, different LME models were specified, and model selection was performed using simulated likelihood ratio tests (with 500 replications for simulation and alpha level set at .05). We performed simulating likelihood ratio tests (as opposed to the likelihood ratio test) as it is recommended when testing for fixed effects and allows comparing arbitrary LME models, i.e., models do not need to be nested and can have different random and fixed effects. The final model was selected that “won” all or most simulated likelihood ratio tests against its competitor models. As conventionally done, winning was determined as *p* < .05 for the more complex model (using the *compare* function in Matlab 2019b). Only models were compared against each other that resulted in a positive definiteness of the Hessian of the objective function with respect to unconstrained parameters at convergence which was used as a criterion to verify optimality of the solution (this results in the different number of competitor models for each research question). The LME formula of the winning models are listed in Table 3 for each research question separately alongside the number of competitor models tested and the number of won competitions.

**Table 3:**
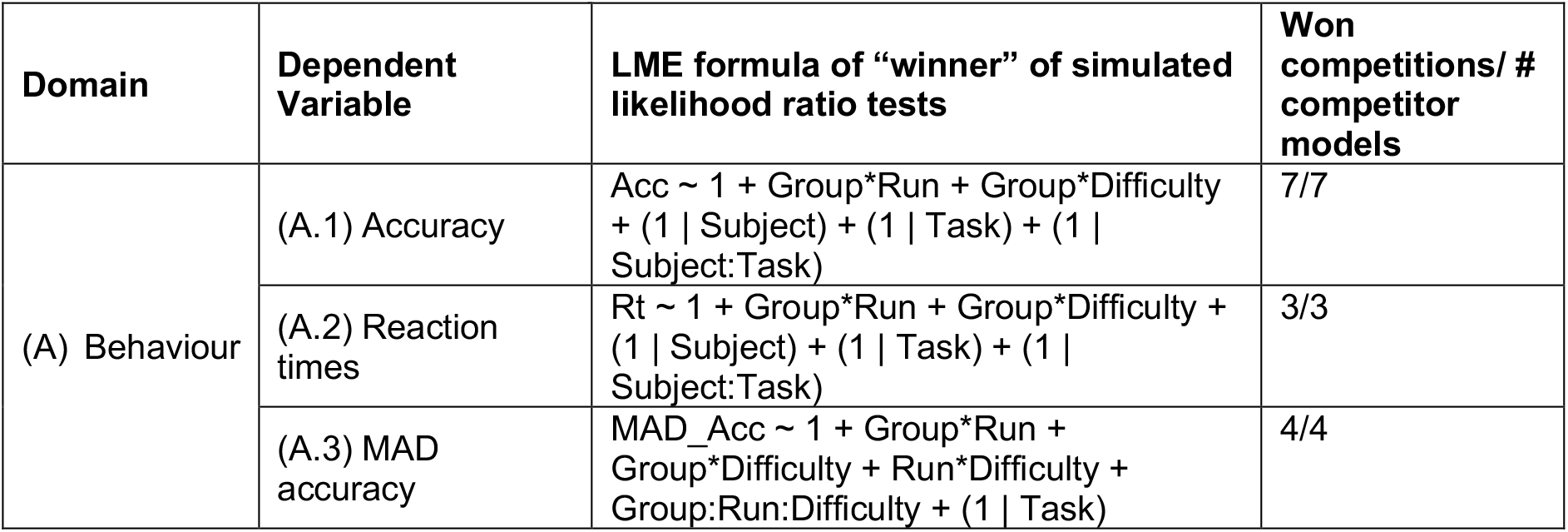
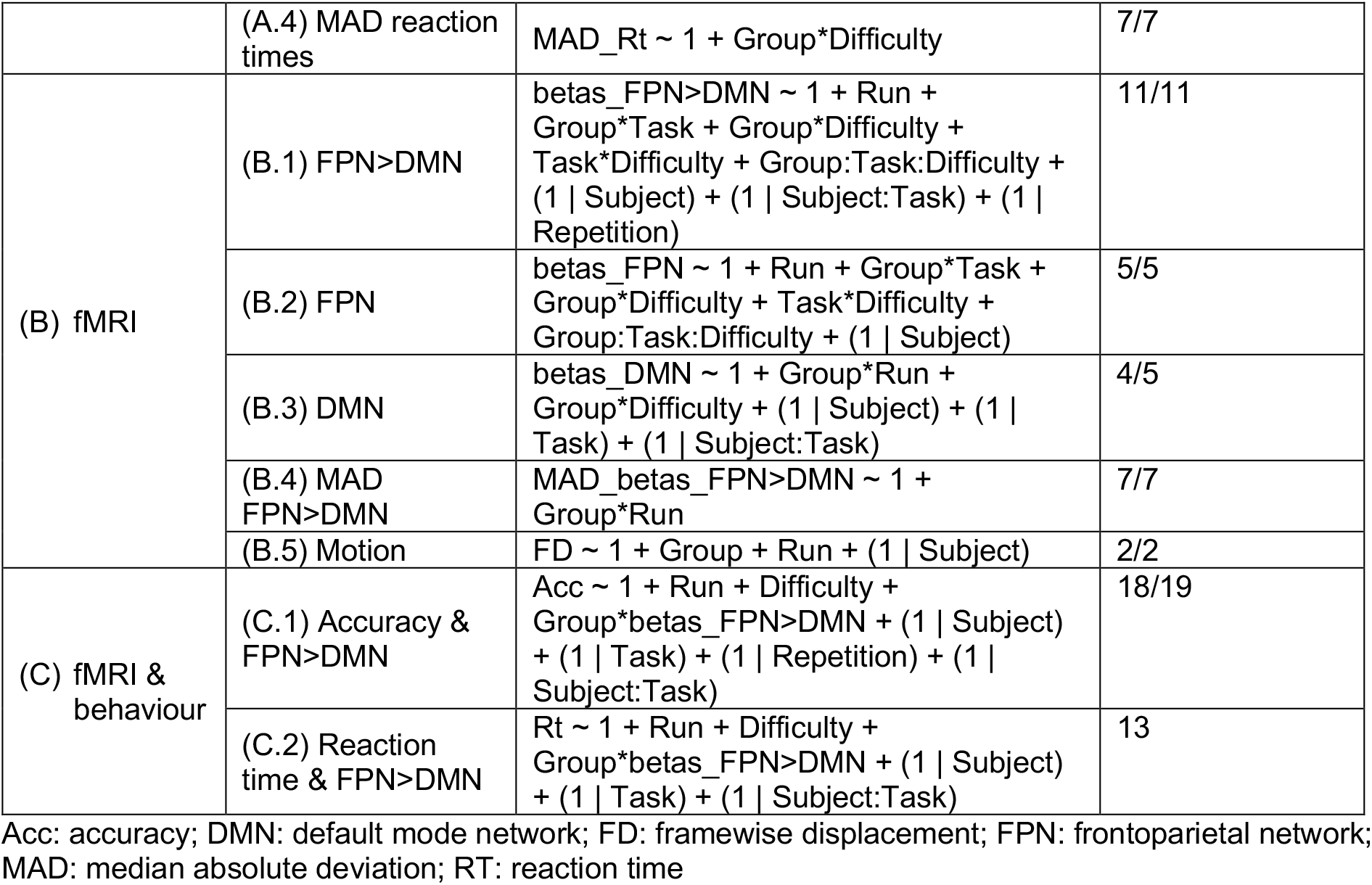
LME formula of winning models for each research question.

### Intra-subject reliability

To assess if the Bayesian optimisation procedure yielded reliable results, all subjects underwent two separate real-time optimisation runs. The Bayesian optimisation algorithm in the second run was thereby completely blind to any data collected in the subject’s previous run or any previous subjects. To assess intra-subject reliability of our results, we computed the spatial correlation (Spearman’s rank correlation) of the Bayesian predictions between the two runs of each subject. For statistical inference, we performed permutation testing (i.e., 10,000 permutations) where we shuffled the FPN>DMN contrast values and corresponding task indices of the second run and then refitted the Gaussian process for these shuffled values before computing the correlation coefficient between the two runs. For each permutation, we then computed the median of the Fisher z-transformed correlation values for each group separately. To correct for multiple comparisons, at each permutation we only kept the maximum of both median values (resembling the “max statistic” method proposed by (Groppe et al., 2011)). The median of our true empirically obtained (Fisher z-transformed) correlation coefficients for patients and controls were then compared to the generated null distribution of maximum median values with a one-sided alpha-level set at .05. Hyper-parameters of the Gaussian process were kept identical to the real-time scenario.

### Assessing dissimilarity of patients’ functional profiles

To assess if patients’ individual profiles were more dissimilar among each other than those of controls, we computed the correlation distance (1- Spearman’s rank correlation (Kriegeskorte, 2008)) of each subject’s functional profile to all other subjects’ individual profiles. For this, we first collapsed the two runs of each subject (i.e., fitting Gaussian process regression on both runs). Next, we computed the correlation distance among all subjects, resulting in a 25 × 25 matrix. From this matrix, we then extracted three different sub-matrices: an 11 × 11 patient-by-patient matrix; a 14 × 14 control-by-control matrix; and an 11 × 14 patient-by-control matrix. For statistical inference, we performed permutation testing (10,000 permutations) where we replicated this procedure but before computing the correlation distance among all subjects, we randomly shuffled the label for patients and controls. We then computed two different t-statistics: (1) for the difference between controls’ dissimilarity (i.e., upper triangle of the symmetrical 14 × 14 matrix) and patients’ dissimilarity (i.e., upper triangle of the 11 × 11 matrix), and (2) for the difference between controls-by-patient dissimilarity (full 11 × 14 matrix) and patients’ dissimilarity (i.e., upper triangle). Finally, our true empirical t-statistics were then compared to the generated null distribution of t-values with a one-sided alpha-level set at .05. To visualise dissimilarity among patients’ functional profiles in 2D, we employed the t-distributed Stochastic Neighbour Embedding (t-SNE) algorithm on the demeaned data (i.e., demeaned by subject) with two different metrics of distance: 1-spearman correlation and Euclidean distance.

### Normalising patient’s FPN-DMN dissociation to control distribution

To assess if there was a single task, for which all patients showed a different FPN-DMN dissociation than controls, we “normalised” each patient’s FPN>DMN contrast value to the control distribution. For normalisation, we computed the modified z-score (Iglewicz & Hoaglin, 1997) *Mi*:

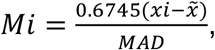

where *xi* refers to the patient’s individual FPN>DMN contrast value, 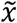 denotes the median of all controls’ FPN>DMN contrast values and *MAD* refers to the median absolute deviation of those values. This analysis was done for each task and difficulty level separately. The normalised z-score is preferred for small samples as the median and MAD are robust measures of central tendency and variance, respectively. A patient’s FPN>DMN contrast value for a particular task was marked significantly different (i.e., “outlier”) when the absolute modified z-score was greater than 1.96. This was a liberal criterion, as commonly a threshold of 3.5 is used.

### Data availability

For Gaussian process regression, we use a Python implementation from: [http://github.com/SheffieldML/GPy]. Python code for the acquisition functions is available from: [http://github.com/romylorenz/AcquisitionFunction]. All relevant data are available from the authors upon reasonable request.

## Acknowledgements

The authors would like to thank the late professor Richard Wise who helped the study take off with his infectious enthusiasm and funding support. Romy Lorenz was funded by the EPSRC (P70597) and is currently funded by the Wellcome Trust (209139/Z/17/Z). Robert Leech received support from the Medical Research Council (MR/R005370/1), the Wellcome/EPSRC Centre for Medical Engineering (WT 203148/Z/16/Z) and the Data to Early Diagnosis and Precision Medicine Industrial Strategy Challenge Fund by the UK Research and Innovation (UKRI) and the National Institute for Health Research (NIHR). Many thanks to Nicolas Langer for immensely helpful statistical guidance and his lab, the University of Zurich as well as Lucie and Sandro Gentile for providing academic and personal refuge for Romy Lorenz in the midst of the Covid-19 pandemic and the sudden closure of US borders.

## Author contributions

FG intellectually conceived the study. RLo, AH, FD, RLe and FG designed the study. FG implemented the experiment under guidance of RLo. RLo, MJ and FG conducted the experiments. RLo analysed the results with support of RLe and FG. MJ provided paragraphs for the Methods section. RLo wrote the manuscript with active involvement from RLe and FG as well as feedback from AH and FD.

